# The xenacoelomorph gonopore is homologous to the bilaterian anus

**DOI:** 10.1101/2025.02.10.637358

**Authors:** Carmen Andrikou, Kevin Pang, Aina Børve, Tsai-Ming Lu, Andreas Hejnol

## Abstract

The bilaterian through gut with an anal opening is a key invention in animals, since it facilitates effective food processing, which allows animals to grow to a larger body size. However, because non-bilaterian animals lack a through gut, the evolution of anus is still debated. The formation of bilaterian hindgut is governed by the spatial expression of several transcription factors (e.g. Caudal and Brachyury) under the control of Wnt signaling. This conserved pattern has been used to support the homology of the anus of protostomes (insects, snails) and deuterostomes (sea urchins, humans). Here we show, that these bilaterian “hindgut” marker genes are expressed around the male gonopore of several xenacoelomorphs, which have a blind gut without an anal opening. These findings suggest a deep evolutionary relationship between the xenacoelomorph male gonopore and the bilaterian anus. Since xenacoelomorphs are the potential sister group to all remaining Bilateria, our results suggest that the bilaterian anus evolved from a male gonopore that came in contact with the digestive endoderm to form the posterior opening.

## Introduction

The evolution of a through gut - a digestive tube with two openings, the mouth and the anus - allowed the unidirectional movement of the food through a regionalized gut and therefore the efficient processing of nutrients. However, a though gut is only found in bilaterians, whilst the closely related Cnidaria possess a blind gut with only one opening (Fig. 1) (Supplementary Fig. 1). How and when the anus emerged during evolution is therefore crucial for understanding the evolution of animal macrofauna and it has been the subject of debates for more than 100 years (Fig. 1) [1-6]. During bilaterian gastrulation the primitive digestive tube compartmentalizes along its anterior-posterior (AP) axis in the distinct sections of foregut, midgut and hindgut. These territories will eventually give rise to the specialized tissues of the adult gut: the foregut will form the esophagus and stomach, the midgut will form the small intestine and the hindgut will form the large intestine and part of the anal canal. The regionalization process of the gut primordia is vital for its proper function and is controlled by the localized expression of a number of transcription factors, several of which are shown to have not only a conserved expression domain but also a conserved pivotal role [7, 8].

**Fig. 1.**
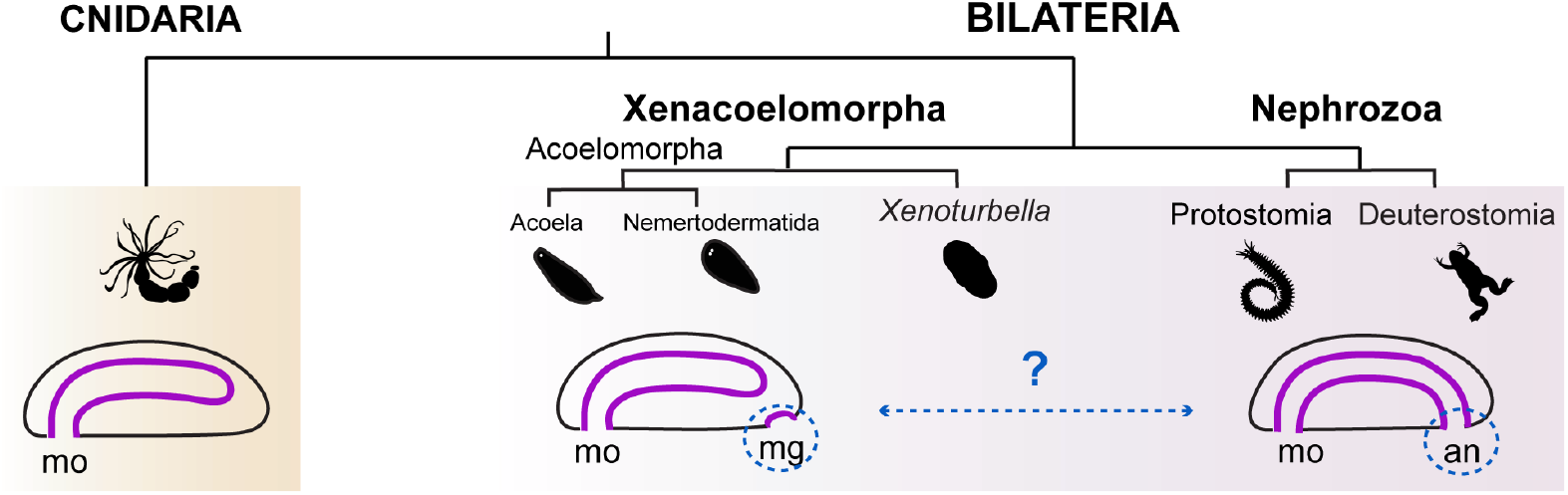
The hypothetical evolutionary transition from a blind gut to a through gut. Cnidarians possess a blind gut with a mouth, while nephrozoans (protostomes and deuterostomes) have a through gut with a mouth and an anus. Xenacoelomorphs, the sister group to the remaining Bilateria, are characterized by a blind gut with a mouth and a male gonopore. Whether the emergence of nephrozoan anus is evolutionary related to the xenacoelomorph male gonopore remains unsolved. an, anus; mg, male gonopore; mo, mouth. Animal illustrations are taken from phylopic.org (CC BY 3.0).

In particular, studies on the molecular patterning of the bilaterian hindgut development, has revealed a set of conserved transcription factors, as well as the Wnt signaling to be expressed in a similar manner among animals [1]. For instance, *caudal* expression is associated to the hindgut formation in members of ecdysozoans [9-11], annelids [12, 13], cephalochordates [14], molluscs [15], echinoderms [16], phoronids [17] and vertebrates [18]. *Brachyury*, on the other hand, can be expressed in both the mouth and the hindgut (e.g., in echinoderms, annelids and molluscs [19-21]), or only the mouth (e.g., in chaetognaths [22]), or only the hindgut (e.g., in ecdysozoans, phoronids and most chordates [17, 23, 24]). Finally, Wnt ligands and receptors are expressed in the hindgut of several animals, often interconnected with *brachyury* and *caudal* [25-35]. This common molecular profile led to the agreement that the hindgut of protostomes and deuterostomes (nephrozoans) share a common ancestry [11, 19].

Xenacoelomorpha (*Xenoturbella* + Acoelomorpha (Nemertodermatida + Acoela)) form the probable sister clade to the remaining bilaterians (Nephrozoa) [36-42], although other positions have been suggested as well [43-45]. Like cnidarians, xenacoelomorphs possess a blind gut with only one opening, the mouth, that corresponds to the bilaterian mouth [2] (Fig. 1) (Supplementary Fig. 1). Their important phylogenetic position therefore holds the key in understanding the evolutionary transition from a blind, sac-like gut to a through gut in bilaterians (Fig. 1). Interestingly, studies on the molecular patterning of the digestive system in two acoel species (*Convolutriloba longifissura* and *Symsagittifera roscoffensis*) showed that *caudal* and *brachyury* [1] are demarcating the region around the male genital opening (gonopore), through which the sperm gets released to the exterior [2, 46]. A putative evolutionary relationship of the acoel male gonopore and the bilaterian hindgut/ anal opening was therefore postulated [2], but not yet thoroughly investigated. Since gonopores do not exist in non-bilaterian animals (Cnidaria, Placozoa and Porifera) (Supplementary Fig. 1), Xenacoelomorpha are therefore important in understanding whether the presence of gonopores is connected to the emergence of a through gut with an anus in the lineage of Bilateria (Fig. 1). To test the hypothesis of the homology of the xenacoelomorph gonopore and bilaterian anus, we investigated more species from the xenacoelomorph clade and revealed the presence of the male gonopore and its relation to the expression of several conserved bilaterian foregut, midgut and hindgut-related genes [1, 26, 47].

## Results

### *Xenoturbella bocki* possesses a male gonopore

In acoelomorphs the male gonopore has been described as part of the ground pattern, since it is present in all acoel and nemertodermatid species investigated so far [48]. In contrast, a female gonopore evolved within the lineage of Acoela and perhaps even more than once [48]. Although the last common ancestor of Acoelomorpha likely possessed only a male gonopore [49], previous studies on individuals of their sister group, *Xenoturbella*, do not describe a male gonopore [50, 51]. We revisited this subject by investigating the species *Xenoturbella bocki*. A single pore with bilaterally arranged white spots was observed posterior-ventrally of the animal, resembling a male gonopore (Fig. 2a-b). Careful examination using immunofluorescence labeling confirmed the presence of two bilateral spermoducts and a central male gonopore at the posterior end of *Xenoturbella bocki* (Fig. 2c). Why the prominent gonopore of *Xenoturbella bocki* has not been described previously remains unclear. Since our specimen was collected during the winter (December), we could perhaps witness a seasonal appearance of the male gonopore. In line with previous works, we could not detect a female gonopore. Therefore, the presence of a male gonopore *Xenoturbella bocki* reflects the condition found in nemertodermatids and acoels. These results suggest that the last common ancestor of Xenacoelomorpha was characterized by a posterior male gonopore and lacked female reproductive openings.

**Fig. 2.**
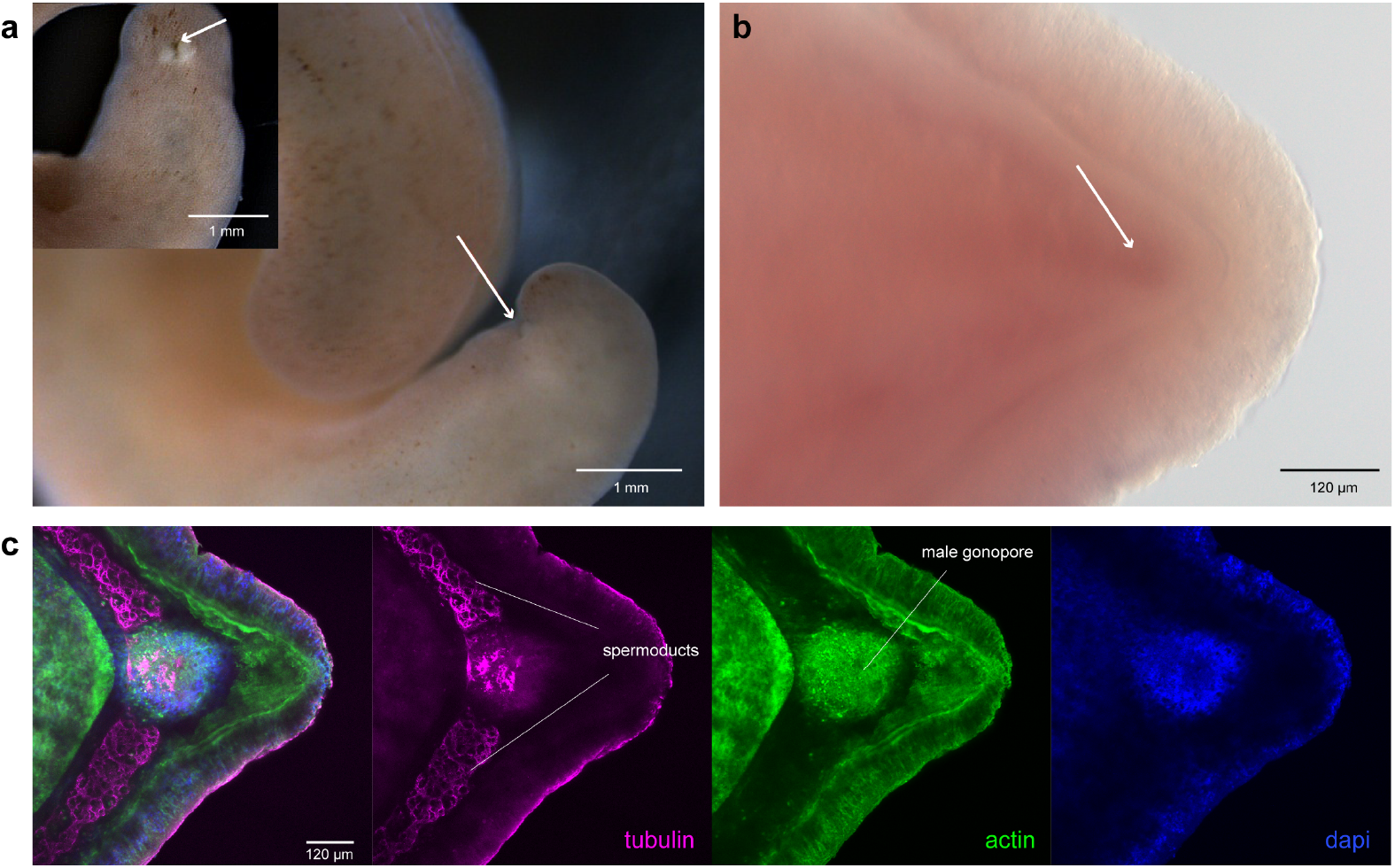
The male gonopore of *Xenoturbella bocki*. **(a, b) Live** image of an adult *Xenoturbella bocki* and indication of a posterior male gonopore (white arrow). **(c)** Immunohistochemistry of anti-tubulin (magenta), anti-actin (green) and nuclei (blue) of the posterior part of a *Xenoturbella bocki* adult. Antitubulin is labeling the sperm. Every fluorescent image is a full projection of merged confocal stacks. Anterior to the left.

### Bilaterian hindgut markers are expressed around the male gonopore of acoels

We first identified orthologous sequences of the foregut/midgut markers *goosecoid (gsc), foxa, gata456, ptfa1* and *hnf4*, the hindgut markers *brachyury, caudal, even skipped (evx)* and *nk2*.*1*, as well as members of the Wnt signal pathway [1, 26, 47] in the available transcriptomes or/ and genomes of *Isodiametra pulchra, Convolutriloba macropyga, Hofstenia miamia* and *Meara stichopi* (Supplementary Fig. 2). We detected some gene duplications within the Acoela for *foxa, gata456, nk2*.*1* and *frizzled 9/10* (Supplementary Fig. 2). Moreover, all acoel Wnt genes grouped in one clade with no clear orthology to other metazoan Wnts, suggesting either independent duplication events within the lineage of acoels or fast evolving sequences, which do not allow their orthologization (Supplementary Fig. 2). On the contrary, orthologs of four nemertodermatid Wnts to other metazoans were found (wnt1, wnt3, wnt5 and wnt11), while 3 nemertodermatid-specific Wnts formed a separate clade (wntx, wnty and wntz) indicating an independent duplication event in nemertodermatids (Supplementary Fig. 2).

We then examined the expression of these genes by Whole Mount *In Situ* Hybridization (WMISH) in adult, sexually matured animals, where the male gonopore was already formed. In *I. pulchra*, the male gonopore is situated at the posterior end of the animal, just posterior to the female gonopore [52] (Fig. 3a). The two paralogs of *foxA* [53] were expressed along the digestive syncytium (Fig. 3aA2), while transcripts of *gata456* [53] were labeling the posterior digestive syncytium and muscles (Fig. 3aA3). Interestingly, the hindgut markers *caudal* (Fig. 3aB1) and *brachyury* (Fig. 3Ab2), but also two members of the Wnt family (*wntb* and *wntg*) (Fig. 3aC2, 7) and their receptors *frizzled9/10a* and *frizzled9/10b* (Supplementary Fig. 3a) were all expressed around the male gonopore. Other members of the Wnt family were expressed in neuronal cells and cells of the posterior tip (Fig. 3aC1, 3-6). Finally, the two paralogs *nk2*.*1a* and *nk2*.*1b* were expressed in the parenchymal muscles and neuronal cells, respectively (Fig. 3aB4). Double WMISH experiments showed that *brachyury* and *caudal* were co-expressed around the male gonopore (Fig. 3a). However, the early juveniles of the acoel *I. pulchra* that don’t have a gonopore yet, were devoid of *brachyury* and *caudal* expression (Supplementary Fig. 4a), suggesting that the expression implies an active role during the male gonopore formation.

**Fig. 3.**
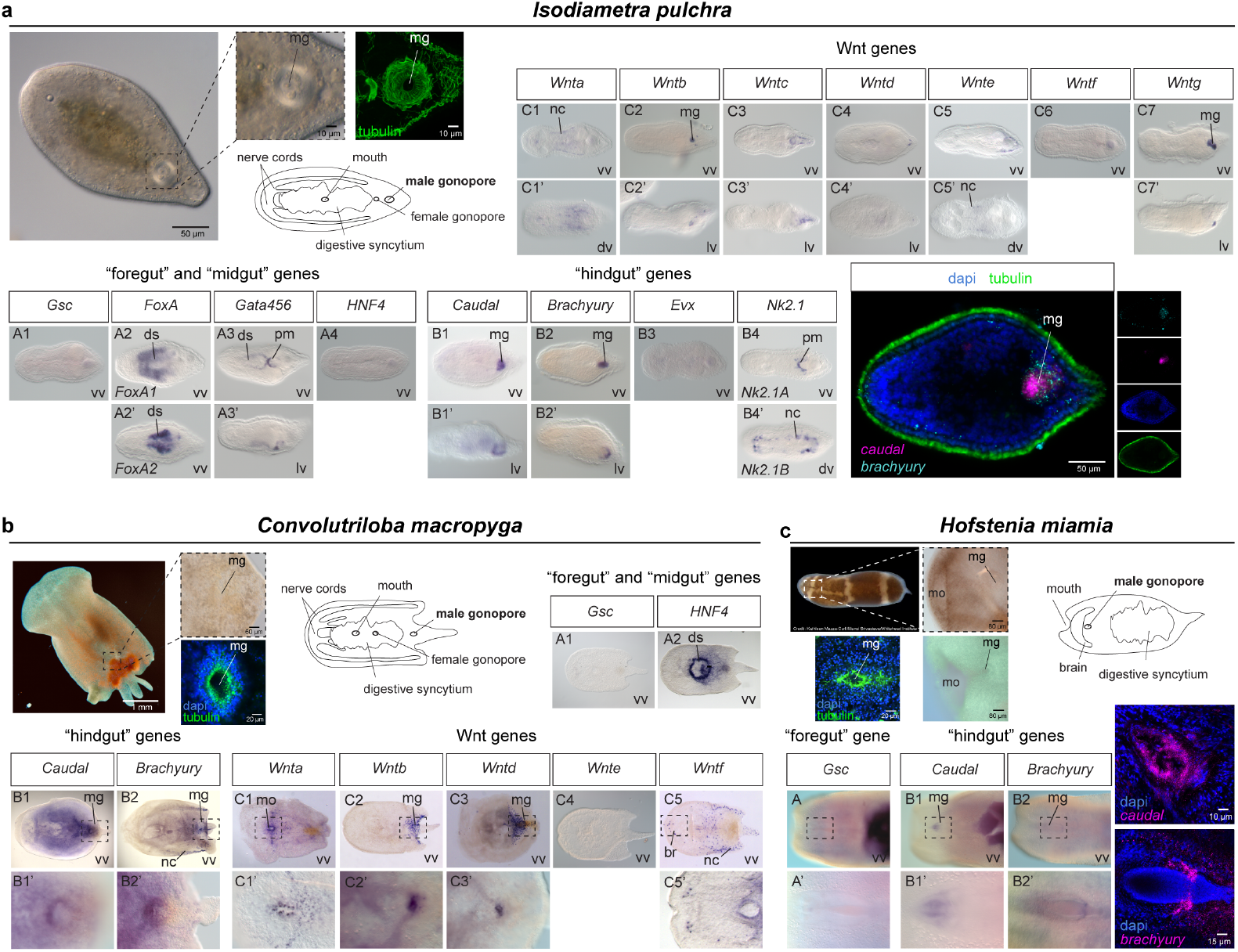
Gene expression of foregut/midgut and hindgut markers in different acoel species. WMISH of the foregut and midgut markers *gsc, foxA, gata456* and *hnf4*, the hindgut markers *caudal* and *brachyury*, and members of the Wnt family in **(a)** *I. pulchra* **(b)** *C. macropyga* and **(c)** *H. miamia*. In *I. pulchra* the male gonopore is depicted with immunofluorescence using anti-actin (green). In fluorescent WMISH *caudal* and *brachyury* expression is in magenta, and nuclei are in blue. In double fluorescent WMISH *brachyury* expression is in cyan, *caudal* is in magenta, tubulin is in green and nuclei are in blue. Drawings are not up to scale. Anterior to the left. ds, digestive syncytium; dv, dorsal view; lv, lateral view; mg, male gonopore; mo, mouth; nc, neural cells; pm, parenchymal muscles; vv, ventral view.

In *C. macropyga*, the male gonopore is located at the posterior region of the animal, posteriorly to the female gonopore [54] (Fig. 3b). Transcripts of *hnf4* were localized in the digestive syncytium (Fig. 3bA2). Again, the hindgut markers *brachyury* and *caudal* (Fig. 3bB1-B2), *wntb* and *wntd* (Fig. 3bC2-3), as well as *frizzled9/10a* (Supplementary Fig. 3b) were expressed around the male gonopore. Other members of the Wnt family were expressed in individual neuronal cells and the mouth (Fig. 3bC1, 5). In immature *C. macropyga* without gonopores, *brachyury* was expressed posteriorly and transcripts of *caudal* were solely localized in the developing nervous system (Supplementary Fig. 4b). The expression of *brachyury* and *caudal in C. macropyga* embryos resembles the previously reported expression in a different *Convolutriloba* species [2]. However, the expression becomes prominent around the male gonopore at the adult stage, suggesting that these genes have a distinct-gonopore specific role. Interestingly, none of the investigated genes was expressed in the female gonopore region in both species. This observation reinforces the plesiomorphic condition of the male gonopore within Acoela and further suggests that the molecular patterning of the two gonopores is different.

### The expression of *brachyury, caudal* and components of the Wnt pathway is gonopore-specific

Since Brachyury, Caudal and the Wnt pathway are involved in the posterior patterning and growth in several animals [27, 31, 55-57], we aimed to discriminate whether the witnessed gene expression is gonopore specific, or related to the overall posterior patterning of the animal. Therefore we investigated the acoel species *H. miamia* where the position of the male gonopore is located anterior in the body, just posterior to the anterior mouth [58] (Fig. 3c). Remarkably, both hindgut markers *brachyury* and *caudal* were expressed around the anterior male gonopore in *H. miamia*, separated from the posterior tip of the animal (Fig. 3cB1-B2). The expression profiles of the Wnt genes (*wnt1*) and the Frizzled receptors (*fz-1)* were reported in a previous publication in the region of the male gonopore [42]. In fact, *wnt1* is an ortholog of *wntg* of *I. pulchra* (Supplementary Fig. 2), which also demarcates the region around the male gonopore (Fig. 3aC7). However, in the early juveniles where the gonopore is not yet formed, *brachyury* was expressed posteriorly and *caudal* was only seen in the developing nervous system, suggesting that their expression around the male gonopore is not a remnant of an earlier embryonic patterning but gonopore-specific (Supplementary Fig. 5). These findings show that in all three investigated acoel species, the expression of the hindgut markers *caudal* and *brachyury*, and members of the Wnt family and Frizzled receptors are consistently expressed around the male gonopore, suggesting that the molecular patterning of the male gonopore is conserved among acoels and bears strong similarities to the bilaterian hindgut.

### Bilaterian hindgut markers are expressed around the male gonopore of nemertodermatids

To understand whether this molecular conservation extends to the lineage of nemertodermatids, we investigated the expression of the aforementioned genes in the nemertodermatid *M. stichopi*. Nemertodermatids are the sister group to acoels, and they possess an epithelial gut versus a digestive syncytium that is an evolutionary derived character of acoels [59]. The male gonopore is placed at the posterior end, while a female gonopore is absent [60] (Fig. 4). *Gsc* was expressed in the mouth (Fig. 4A1), and transcripts of *foxa* (Fig. 4A2) and *hnf4* (Fig. 4A4) were localized in the entire gut. *Ptfa1* was expressed only in the posterior part of the gut (Fig. 4A5). The hindgut markers *caudal* (Fig. 4B1) and *brachyury* (Fig. 4B2) were expressed around the male gonopore, while *evx* was expressed at the posterior tip (Fig. 4B3) and *nk2*.*1* showed a ventral expression (Fig. 4B4). Moreover, three members of the Wnt family (*wnt1, wnty, wntz*) (Fig. 4C1, C6-7), and the receptor *frizzled9/10* (Supplementary Fig. 3c) were expressed around the male gonopore. Other members of the Wnt family were marking neuronal cells and cells of the posterior tip (Fig. 4C2-5). Transcripts of *nk2*.*1* were additionally demarcating the female gonadal region (Fig. 4B4) and *brachyury* was also expressed in the mouth (Fig. 4B2). Double WMISH analysis confirmed that *brachyury* and *caudal* were co-expressed around the male gonopore and the same was true for *brachyury* and *wnt1* (Fig. 4). Finally, similar to the case in the acoels, the immature juveniles of the nemertodermatid *M. stichopi* that do not possess a gonopore yet, *brachyury* expression was restricted in the future mouth region -the mouth forms later in nemertodermatids [61]- and *caudal* was absent (Supplementary Fig. 6).

**Fig. 4.**
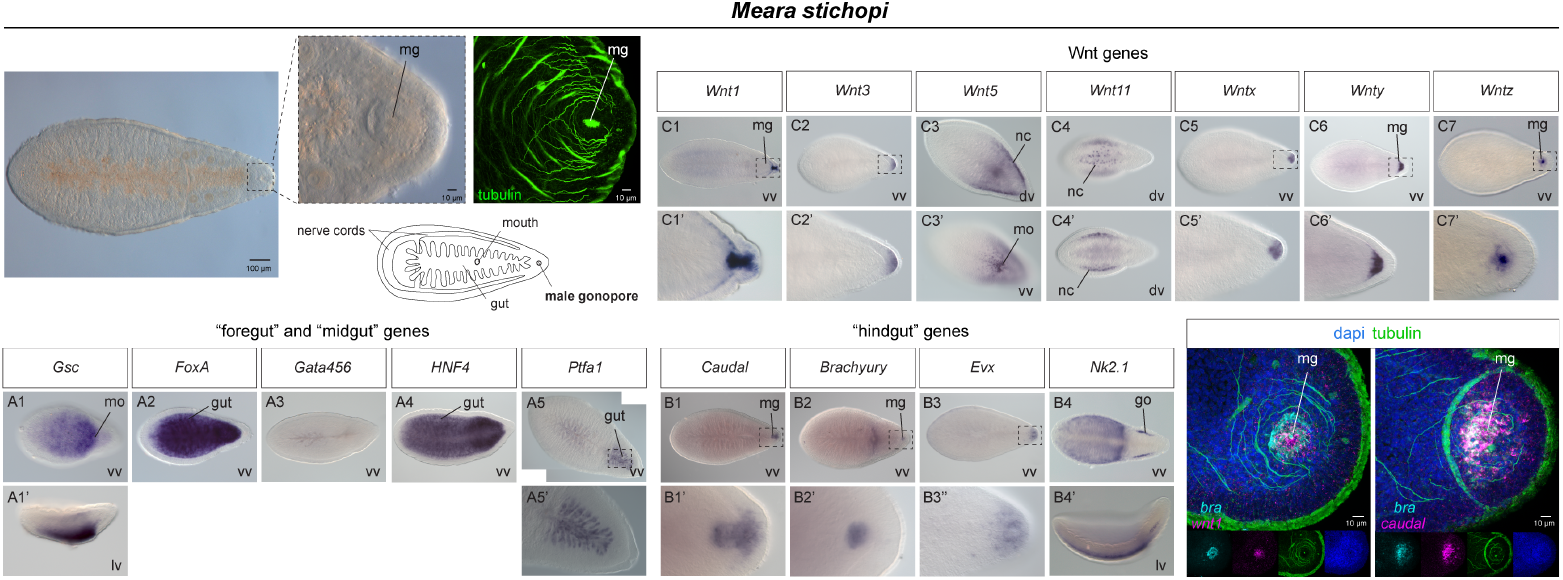
Gene expression of foregut/ midgut and hindgut markers in nemertodermatids. WMISH of the foregut and midgut markers *gsc, foxA, gata456, ptf1* and *hnf4*, the hindgut markers *caudal* and *brachyury*, and members of the Wnt family in *M. stichopi*. The male gonopore is depicted with immunofluorescence using anti-tubulin (green). In double fluorescent WMISH *brachyury* expression is in cyan, *caudal* and *wnt1* expression is in magenta, tubulin is in green and nuclei are in blue. Drawings are not up to scale. Anterior to the left. dv, dorsal view; go, female gonads; lv, lateral view; mg, male gonopore; mo, mouth; nc, neural cells; vv, ventral view.

In general, these findings demonstrate that in all investigated acoel and nemertodermatid species, the bilaterian hindgut markers are specifically expressed in the male gonopore region during its formation. However, the same is not true for the foregut/midgut markers, which are only expressed in regions associated with the digestive system of acoelomorphs. Therefore, these results support the homology of the male gonopore and the hindgut [2].

## Discussion

### An ancestral molecular patterning is utilized for hindgut and male gonopore formation

The hindgut of most bilaterians forms by the expression of *brachyury* and *caudal* transcriptional regulators, as well as the Wnt signaling pathway [1]. Xenacoelomorpha, the sister group to all remaining bilaterians [36-42], lacks a through gut but acoels and nemertodermatids use these bilaterian hindgut–related genes and the Wnt signaling cascade, in patterning their male gonopore (Fig. 5a). In non-bilaterians however, that also lack a through gut, the expression of these genes is rather broad and not assigned to a common developmental or morphogenetic process. For instance, *brachyury* seems to exhibit a variety of expression patterns, such as in the stomodeal/pharyngeal cells of the Ctenophora [62], the choanocytes of the Porifera [63], the opening of the digestive system in cnidarians [64, 65], and isolated cells located at the edge of the organism in Placozoa [66]. In contrast, *caudal* gene appears to be missing from the Ctenophora, Placozoa and Porifera [67]. In cnidarians a *caudal /lox* ortholog is expressed differently among species (e.g., in the ventral pair of mesenteries of *Nematostella* [68] and in both the oral and aboral pole in *Clytia* [69]). Finally, regarding the Wnt signaling, it is believed that in non-bilaterians was mainly necessary for the formation of a primary, anterior–posterior body axis (a function also witnessed in Bilateria) [70]. The absence of a clear *caudal* ortholog, the variability of *brachyury* expression and the role of wnt pathway as a polarization signal in non-bilaterian animals imply that the assignment of these genes in patterning the posterior digestive system (hindgut) and the xenacoelomorph gonopore likely occurred after the Bilateria/ Cnidaria split. The male gonopore of most xenacoelomorphs forms by involution of the posterior ectoderm, a similar process that occurs during the anus formation of several metazoans [3, 5], suggesting that the same ancestral posterior ectodermal molecular patterning is utilized for both the hindgut/anal canal and male gonopore formation. Therefore, since the male gonopore is part of the xenacoelomorph ground pattern, we propose that the male gonopore of xenacoelomorphs and the bilaterian anus are homologous structures.

**Fig. 5.**
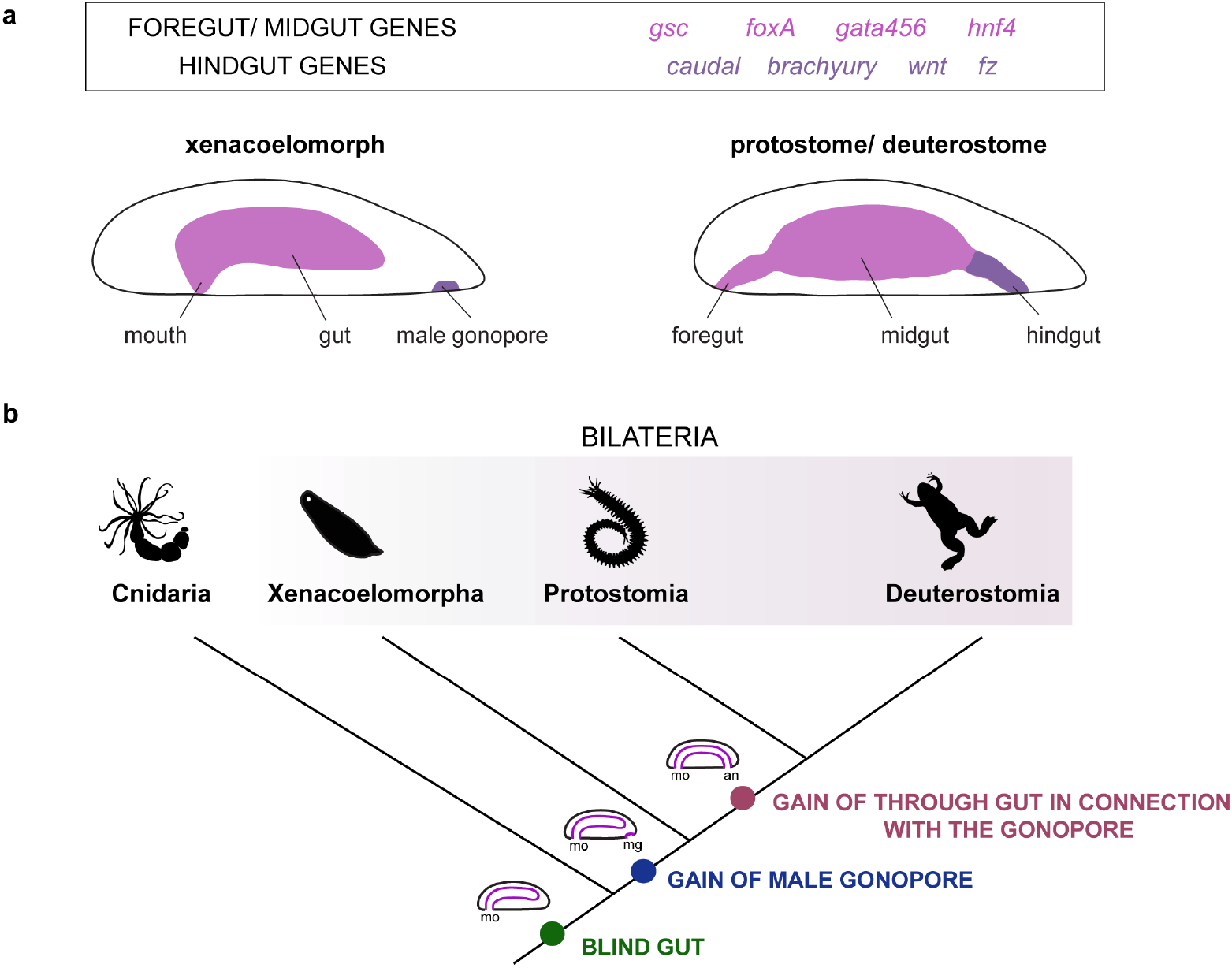
Homology of the xenacoelomorph male gonopore and the bilaterian anus. **(a)** Summary of bilaterian foregut/ midgut and hindgut–related gene expression in Nephrozoa (protostomes+deuterostomes) and Acoelomorpha. *Brachyury, caudal* and *wnt* ligands and receptors are expressed in the male gonopore of acoelomorphs and the hindgut of nephrozoans, suggesting their homology. The hindgut markers were initially expressed in the gonopore and afterwards they were assigned in patterning the hindgut/ anal opening. **(b)** The last common bilaterian ancestor possessed a blind gut and a gonopore, and gained a hindgut/ anal opening in connection with the gonopore after the Xenacoelomopha/ Nephrozoa split.

### The bilaterian anus evolved from a male gonopore

The direction of evolutionary transition between a gonopore and a hindgut with an anus can be interpreted in different ways, depending on whether the absence of an anal opening is plesiomorphic for the Bilateria. Morphologists have developed different hypotheses in the past, where Platyhelminthes and Acoelomorpha were selected as examples of transition states due to their variable digestive and genital system architectures. The main difference was whether the last common ancestor possessed a gonopore and a blind gut without an anal opening [71, 72] or a through gut with a cloaca combining both genital and digestive functions [73]. The presence of cloaca within animals as well as the gonopore-oral fusion witnessed in species of Platyhelminthes (Supplementary Fig. 1), suggests that a connection between the digestive and the reproductive system is either easy to evolve convergently or shares a common ancestry.

Revisiting the subject adding molecular data and using better phylogenies we can now propose that the last common bilaterian ancestor likely possessed a blind digestive tract with an mid-ventrally placed single opening that functioned as a mouth expressing *gsc, foxa, gata456* and *hfn4*, and a posterior male gonopore expressing *caudal, brachyury* formed under a Wnt signal, similar to what is found in acoels and nemertodermatids. This (ectodermal) male gonopore region was later connected with the posterior domain of the endodermal blind digestive tract resulting in the acquisition of a hindgut and the subsequent formation of an anus, perhaps in the form of a cloaca (Fig. 5b). These “gonopore” genes were then recruited in patterning the hindgut, suggesting that the anus evolved in connection with the male gonopore. The alternative but less parsimonious scenario would be that the last common bilaterian ancestor possessed a through gut with an anus expressing hindgut-related genes, and the hindgut with the anal opening was secondarily lost in the lineage of xenacoelomorphs. The expression of “hindgut” markers around the male gonopore of acoelomorphs should be then interpreted as a secondary recruitment of these genes to the posterior ectodermal–endodermal boundary, where the male gonopore forms. However, although an ancestral bilaterian condition of a through gut cannot be excluded, a plesiomorphic blind gut that gained later an anal opening in connection with the gonopore seems more likely (Fig. 5b). In fact, these genes are never expressed in association with the digestive system in acoelomorphs indicating that the posterior portion of the gut was not evolved yet in this lineage.

Our findings provide strong molecular evidence for the homology of the male gonopore of the Xenacoelomorpha and the anus of the Bilateria.

## Materials and methods

### Animals

Adult specimens of *Isodiametra pulchra* Smith and Bush, 1991, *Meara stichopi* Westblad, 1949, *Hofstenia miamia* Correa, 1960 and *Convolutriloba macropyga*, Shannon and Achatz, 2007 were kept and handled as previously described [54, 74, 75]. *Xenoturbella bocki*, Westblad, 1949 were obtained from Gullmarsfjord, Sweden.

### Gene cloning and orthology assignment

Putative orthologous sequences of genes of interest were identified by tBLASTx search against the transcriptome (SRR2681926) of *I. pulchra*, the transcriptome (SRR2681155) and draft genome of *M. stichopi*, the transcriptome (SRX1343815) of *C. macropyga* and the transcriptome (PRJNA241459) and genome (GCA_004352715) of *H. miamia*. Gene orthology of genes of interest identified by tBLASTx was tested by reciprocal BLAST against NCBI Genbank and followed by phylogenetic analyses. Amino acid alignments were made with MAFFT, followed by automated alignment trimming with trimAl. RAxML-NG (version 8.2.9) was used to conduct a maximum likelihood phylogenetic analysis. Fragments of the genes of interest were amplified from cDNA of each animal by PCR using gene specific primers. PCR products were purified and cloned into a pGEM-T Easy vector (Promega, USA) according to the manufacturer’ s instruction and the identity of inserts confirmed by sequencing.

### Whole Mount *In Situ* Hybridization

Embryos were manually collected, fixed and processed for colorimetric and double fluorescent *In situ* hybridization as described in [2, 76]. Labeled antisense RNA probes were transcribed from linearized DNA using digoxigenin-11-UTP (Roche, USA) or labelled with DNP (Mirus, USA) according to the manufacturer’s instructions.

### Whole Mount Immunohistochemistry

Animals were collected manually, fixed in 4% paraformaldehyde in SW for 60 minutes, washed 3 times in PBT and incubated in 4% sheep serum in PBT for 30 min. The animals were then incubated with commercially available primary antibodies (anti-acetylated tubulin mouse monoclonal antibody, dilution 1:250 (Sigma-Aldrich, USA) and anti-actin mouse monoclonal antibody, dilution 1:400 (Seven Hills Bioreagents) overnight at 4°C, washed 5 times in PBT, and followed by incubation in 4% sheep serum in PBT for 30 min. Specimens were then incubated with a secondary antibody overnight at 4°C followed by 5 washes in PTW. Nuclei were stained with DAPI (Invitrogen).

### Documentation

Colorimetric WMISH specimens were imaged with a Zeiss AxioCam HRc mounted on a Zeiss Axioscope A1 equipped with Nomarski optics and processed through Photoshop CS6 (Adobe). Fluorescent-labeled specimens were analyzed with a SP5 confocal laser microscope (Leica, Germany) and processed by the ImageJ software version 2.0.0-rc-42/1.50d (Wayne Rasband, NIH). Figure plates were arranged with Illustrator CS6 (Adobe).

## Notes

### Competing Interest Statement

The authors have declared no competing interest.

