## Supplemental for "The xenacoelomorph gonopore is homologous to the bilaterian anus"

### Supplement

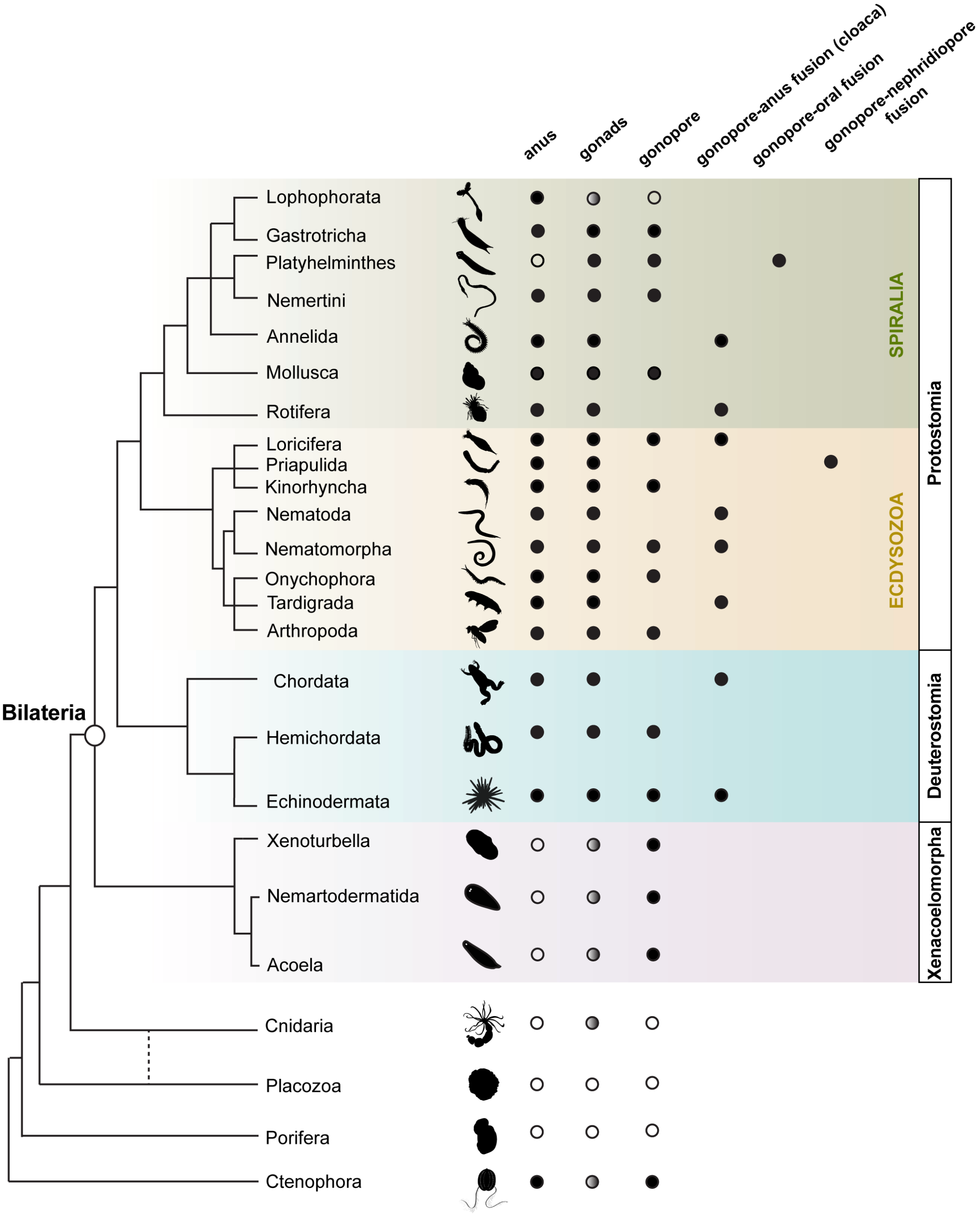


**Supplementary Fig. 1 Phylogenetic placement of the anus, gonopore, gonads and cloaca.** Blind digestive systems (gut without anal opening) are found in non-bilaterians, Xenacoelomorphs and Platyhelminthes. Gonopores are found in members of all animal groups, while cloaca is only witnessed in members of protostomes and deuterostomes. Gonopores can vary in their location and morphology; they can be placed posterior-ventrally (e.g., in arthropods [1, 2]), ventro-laterally (e.g., in *nemerteans [3])*, anterior-ventrally (e.g., in *Hofsteniida* (acoelomorphs) [4]), or even dorsally (e.g., in Catenulida (platyhelminthes) [5, 6]). Moreover, gonopores can exist either as separate, sometimes even segmental, entities (e.g., in nemerteans, arthropods and acoelomorphs) [1-3, 7], or in connection to either the posterior portion of the digestive system (anus) forming a common opening called cloaca (e.g., in chordates, loriciferans, rotifers, holothurians, nematomorphs and tardigrades) [8-14] or the anterior portion of the digestive system (mouth) (e.g., in Lecithoepitheliata and Prolecithophora (Platyhelminthes) [5, 6]. Non bilaterians lack gonopores and they release their gametes either via the aquaferous system (e.g., in Porifera [15]) or through the mouth opening (e.g., in Cnidaria [16]). White circles show the absence of gonads and grey circles indicate that epithelial lined gonads are not present in these species. Animal illustrations are taken from phylopic.org (CC BY 3.0).

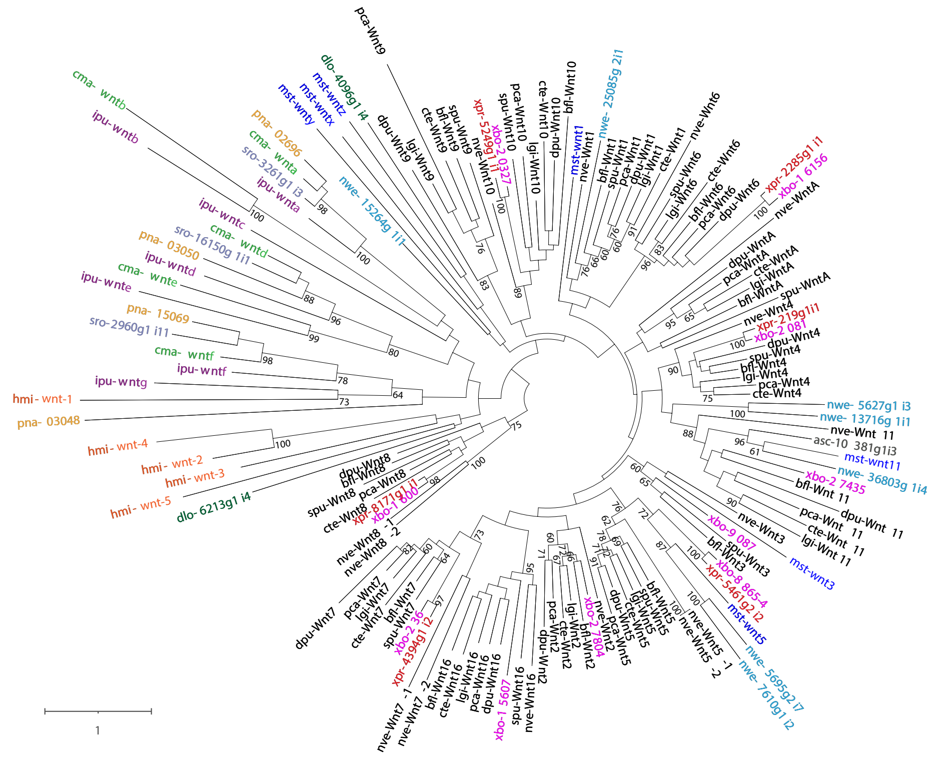

**Supplementary Fig. 2 Wnt gene orthology analysis.** The phylogenetic analysis of Wnt sequences. The alignment matrix used for wnt tree reconstruction was generated using MAFFT, followed by automated alignment trimming with trimAl. The tree was reconstructed using RAxML-NG with a maximum-likelihood optimality criterion. Names of genes or proteins, if available, follow the name of organism(s). *Isodiametra pulchra*, *Meara stichopi*, *Hofstenia miamia* and *Convolutriloba macropyga* sequences are highlighted in purple, blue, orange and light green, respectively. Other xenacoelomorph sequences included in the analysis are taken from the transcriptomes of *Xenoturbella bocki* (in pink), *Xenoturbella profunda* (in red), *Nemertoderma westbladi* (in cyan), *Praesagittifera naikaiensis* (in yellow), *Symsagittifera roscoffensis* (in grey), *Diopisthoporus longitubus* (in dark green) and *Ascoparia sp.* (in grey).

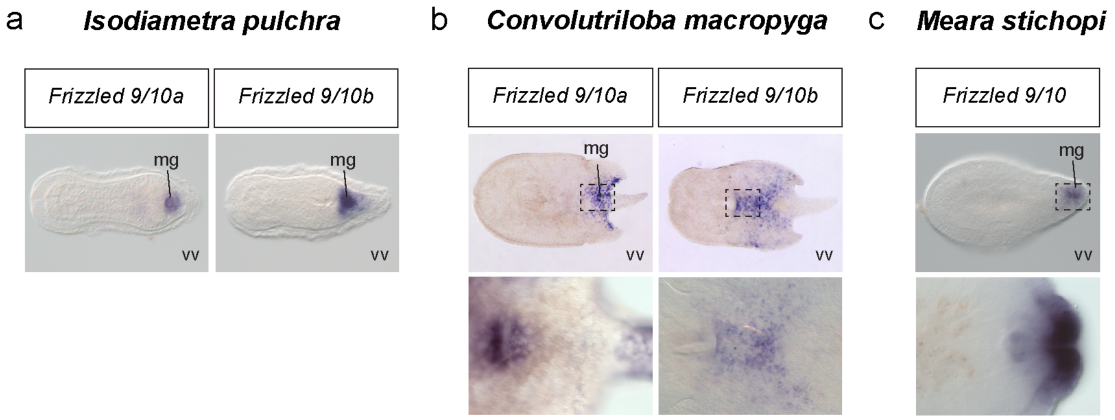


**Supplementary Fig. 3 WMISH of *frizzled9/10* genes in (a) *Isodiametra pulchra*, (b) *Meara stichopi* and (c) *Convolutriloba macropyga.*** Anterior to the left. mg, male gonopore; vv, ventral view.

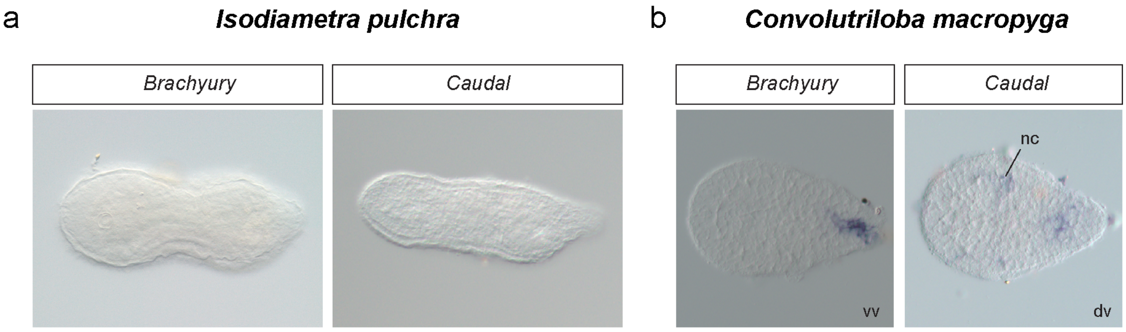


**Supplementary Fig. 4 WMISH of *brachyury* and *caudal* in (a) *Isodiametra pulchra* and (b) *Convolutriloba macropyga* juveniles.** Anterior to the left. dv, dorsal view; nc, neural cells; vv, ventral view.

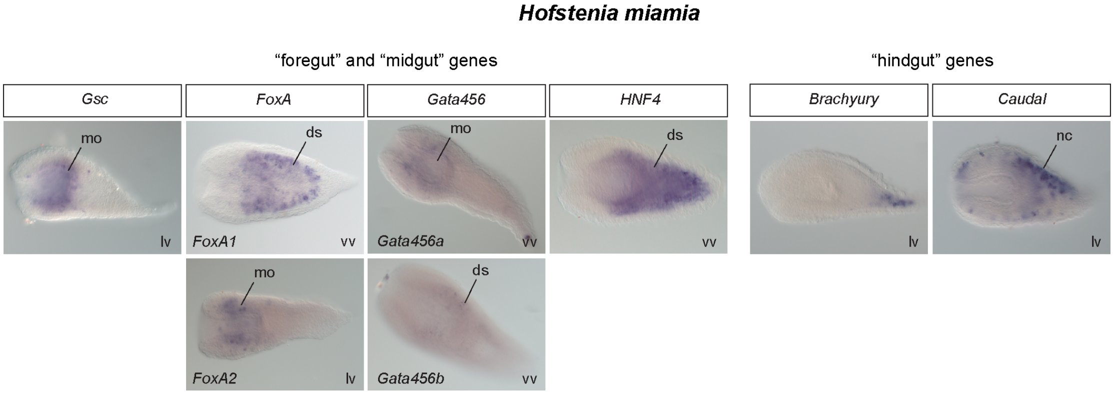


**Supplementary Fig. 5 WMISH of foregut/ midgut markers and hindgut markers in *Hofstenia miamia* juveniles.** WMISH of the foregut and midgut markers *gsc*, *foxA, gata456* and *hnf4* and the hindgut markers *caudal* and *brachyury* in *Hofstenia miamia* juveniles. Anterior to the left. ds, digestive syncytium; lv, lateral view; mo, mouth; nc, neural cells; vv, ventral view.

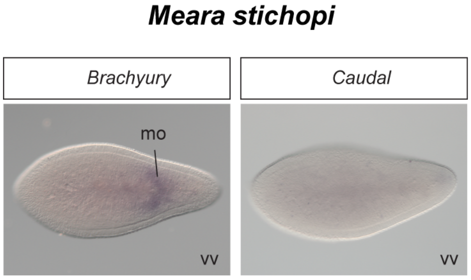


**Supplementary Fig. 6 WMISH of *brachyury* and *caudal* in *Meara stichopi* juveniles.** Anterior to the left. mo, mouth; vv, ventral view.
